# Evidence, evolution and pattern of contractile elements in villous vascular walls and stroma of human placentas of premature infants of different gestational ages. An immuno-morphologic comparative study

**DOI:** 10.1101/2021.01.05.425374

**Authors:** Valerio Gaetano Vellone, Michele Paudice, Rita Bianchi, Giulia Scaglione, Chiara Maria Biatta, Davide Buffi, Francesca Buffelli, Fabio Barra, Simone Ferrero, Ezio Fulcheri

## Abstract

The contractile elements of the human placenta villous tree represent a topic of interest and many issues persist still open. Histology of the stroma and muscular wall and their evolution, in relation with the gestational age, remains to be clarified for a deeper understanding of the adaptive potential and pathogenetic mechanisms.

In our study, 56 premature placentas (21-36 wks) were considered, sub-divided into four groups based on age of gestation and compared to 23 at-term placentas (37-40 wks). All cases were tested with anti-smooth muscle actin (SMA) and anti-desmin antibodies to identify the contractile elements in the stroma and in vascular walls of villi.

SMA and desmin staining show evident decreased expressions during the pregnancy (temporal variation) and from proximal to distal part of the villous tree (spatial variation) being higher in the stem villi.

Both pre-term and at-term placentas showed persisting, although variable, positivity for SMA and desmin staining in the stroma and in the vessel walls of the mature intermediate and terminal villi. This represents an unexpected finding and nothing alike has been previously reported in literature.

Both highly premature and term placentas seem to maintain contractile components within each type of villi, represented by both myofibroblasts and mature smooth muscular cells. These components may be present in both villous vascular walls and stroma, albeit with different staining intensity.

This finding allows us to imagine an active function in the regulation of the blood flow, not only in stem and intermediate immature villi but even in smaller villi.

## 1. Introduction

The histopathological examination of the placenta is a unique opportunity to assess the state of health of both the mother and the foetus. The microanatomy of the stroma and blood vessels wall of the villous tree represent a topic of interest but many issues remain still open. Histology and the evolution of the contractile elements of stroma and muscular wall of the vessels should be clarified in relation with the gestational age and possible adaptive phenomena. Histological and immunophenotypic features of villi stroma and blood vessel in full term placentas appear well consolidated in current literature [1-8].

In Kaufmann’s studies [9] a distinction between reticular and perivascular stroma of stem villi of term placentas is attempted [3,9], and the absence of leiomuscular contractile component is reported in the reticular stroma of immature intermediate villi and in the superficial cellular rim of stem villi [2,3,9]. It is also stated that there is no leiomuscular component in mature intermediate and terminal villi [3,9]. These observations contrast sharply with the characteristics of first trimester placentas, in which the stromal contractile component is ubiquitously present in all degrees of villous branching [3]. Furthermore, previous studies lack of precise references on the vascular walls of villi, while the attention is often focused on composition of the reticular and perivascular stroma of villi [2,3,7,9,21,22]. The evolution of the characteristics of villi between I° and III° trimester is as interesting as little known.

Another point of concernment is represented by the decrease, till to absence, during gestation of the contractile component of the blood vessel walls of mature intermediate and terminal villi [3].

The presented paper is focused on identifying the contractile components in both the stroma and the vascular walls of the premature placentas’ villi in different gestational ages, and on the comparison with the contractile components of term placentas to clarify their evolution.

## 2. Materials and methods

The reported study is retrospectively based on the computerized database of the Fetal and Perinatal Pathology Unit of the Giannina Gaslini Children’s Hospital in Genoa.

According to Vohr and Stephens [10,11,12], we considered as pre-term placentas expelled before completing the 37th gestational week and as highly premature if expelled with a gestational age between the 23rd and the 33rd weeks. A total of 56 preterm placentas were selected for the study. Although the development of the villous tree is continuous, in our study highly premature placentas were divided into four subgroups on the base of the progressive appearance of terminal villi: 21-24 weeks, 25-28 weeks, 29-32 weeks and 33-36 weeks; as a control group, 23 randomly selected placentas between 37th and 40th week of gestation were included.

All the specimens were extensively sampled, routinely processed and embedded in paraffin. Additional slides have been cut from the most significant block and tested with avidin-biotin-streptokinase immunohistochemistry (IHC) method, using the Benchmark XT, Ventana Medical System, Tucson Arizona, USA. All the chorionic disc sections have been tested for Smooth Muscle Actin (SMA) (Novocastra, clone HHF35, IgG1, 1:50 dilution) and Desmin (Novocastra clone De-R-11, IgG1 class, 1:100 dilution).

We defined the parameters of adequate maturation among gestational age groups, and divided premature cases into four ages in accordance with the following criteria: villous ramification description, foetal-maternal exchange elements differentiation, features of terminal villi, trophoblastic replicative activity, features of villi blood vessels wall, and features of villi stroma [1,9,13]. Intensity of IHC staining was graded as follow: no staining (neg), weak (-/+), mild (+), moderate (++), and strong (+++). All the cases were examined by expert pathologists in the field of fetal and perinatal pathology (EF, VGV).

Our study, which has been authorized by the Local Ethics Committee, is completely retrospective and based on computerized archive. The extrapolated data from the histological examination and the clinical history were entered on an anonymous MS Excel© spreadsheet and compared.

## 3. Results

Description of villous ramification features in each gestational group is shown in table 1. The staining intensities for SMA and desmin, according to the gestational age and distinguishing between villi vessels and stroma, are illustrated in figures 1 and 2. As general observation, SMA highlights more elements and cellular types, while desmin highlights less elements but has a stronger staining.

**Table 1:**
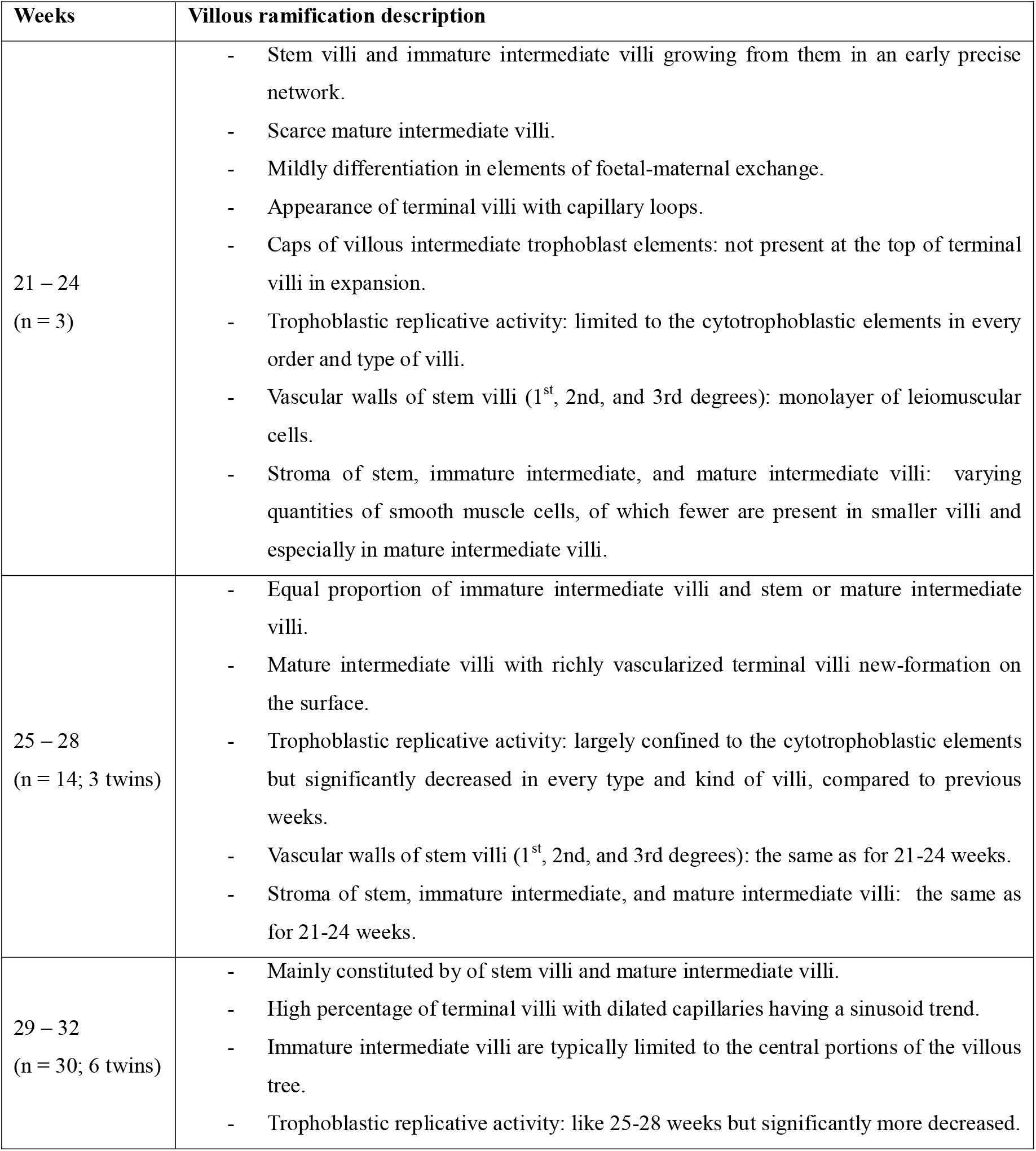
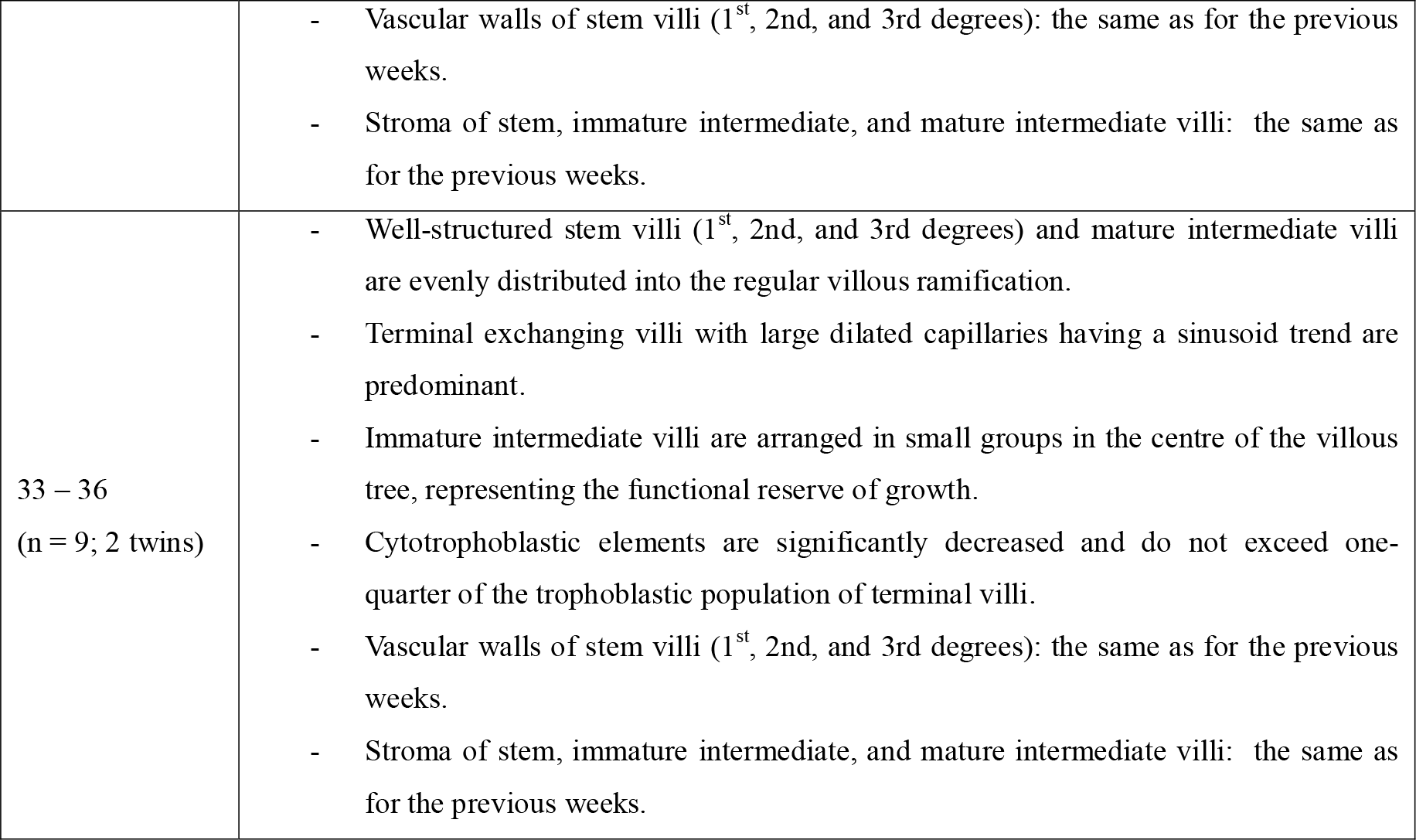
Villous ramification description in the gestational age categories

**Figure 1:**
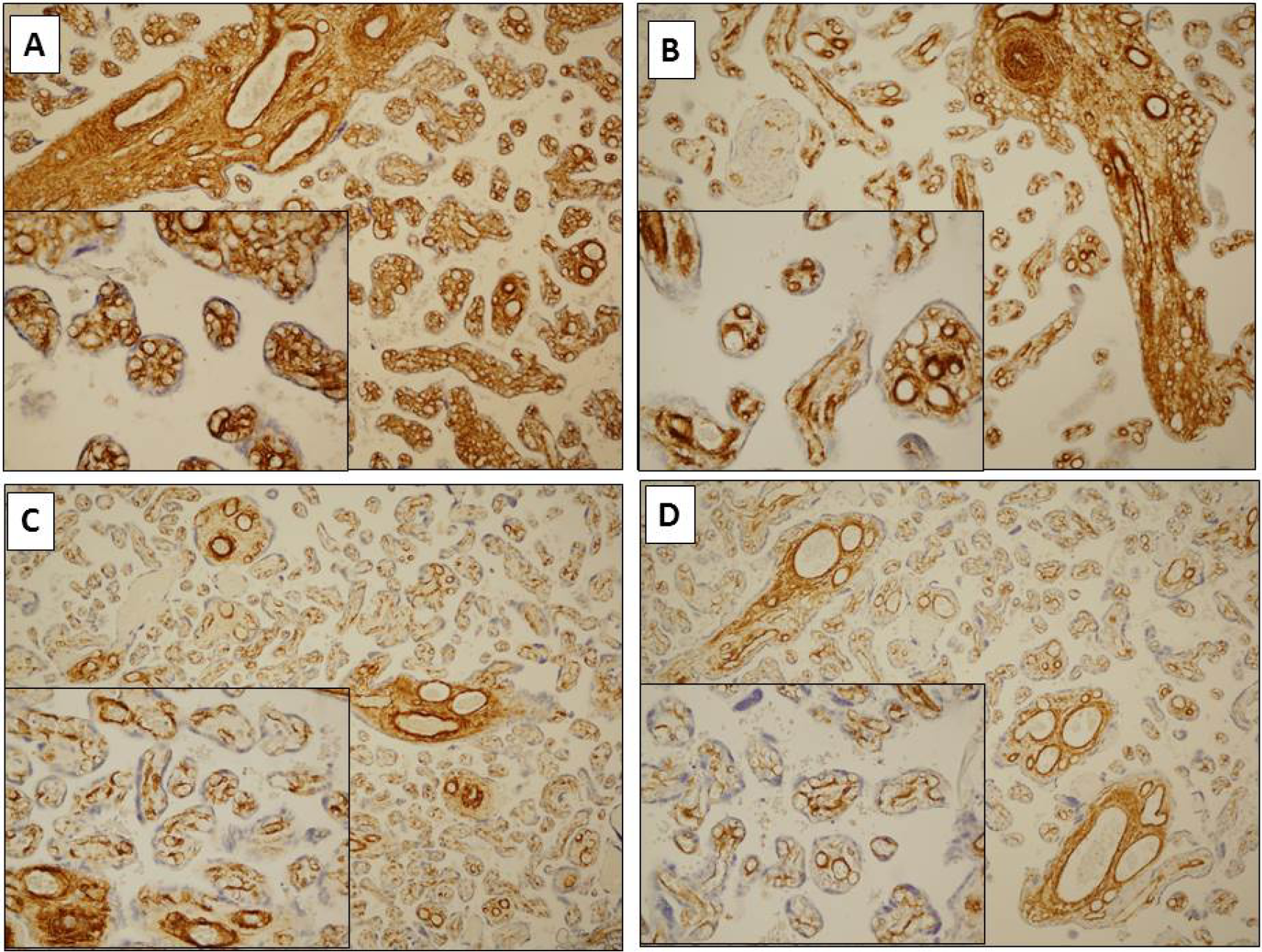
Smooth muscle actin (SMA) Immunohistochemistry. A) Weeks 21-24 (IHC 100x; lower left 400x): moderate to strong stain in the vessel walls and mild to moderate stain in the stroma of stem villi and immature intermediate villi. Moderate stain in the vessel walls and a mild to moderate stain in the stroma of intermediate mature villi. B) Weeks 25-28 (IHC 100x; lower left 400x): mild to moderate stain in the vessel walls and moderate to strong stain in the stroma of stem villi and intermediate immature villi; variable (from negative to strong) stain in the vessel walls and weak to moderate stain in the stroma of mature intermediate villi. Variable (from negative to strong) stain in the vessel walls and a variable (from weak to moderate) stain in the stroma of terminal villi. C) Weeks 29-32 (IHC 100x; lower left 400x): moderate to strong stain in the vessel walls and mild to moderate stain in the stroma of stem villi and intermediate immature villi. Variable (from weak to strong) stain in the vessel walls and weak to moderate in the stroma of intermediate mature villi; variable (from negative to strong) stain in the vessel walls and a variable (from negative to moderate) stain in the stroma of terminal villi. D) Weeks 33-36 (IHC 100x; lower left 400x): moderate to strong stain in the vessel walls and mild to moderate stain in the stroma of stem villi; moderate to strong stain in the vessel walls and mild to moderate stain in the stroma of rare immature intermediate villi. Mild to moderate stain in the vessel walls and in the stroma of intermediate mature villi; variable (from weak to moderate) stain in the vessel walls and variable (from negative to moderate) stain in the stroma of terminal villi.

**Figure 2:**
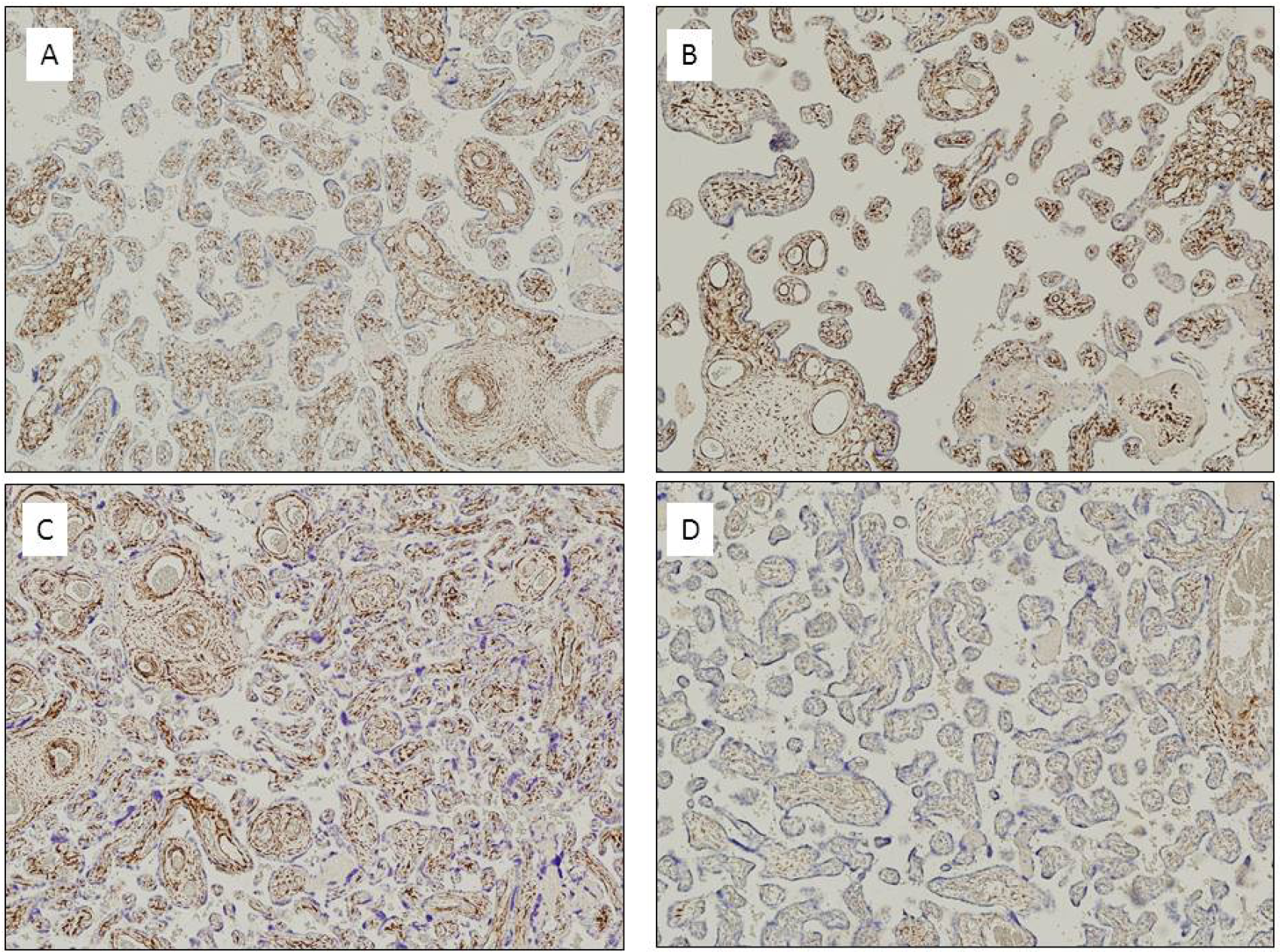
Desmin Immunohistochemistry. A) Weeks 21-24 (IHC 100X): moderate stain in the vessel walls and strong stain in the stroma of stem villi and immature intermediate villi. Mild stain in the vessel walls and a mild to moderate stain in the stroma of intermediate mature villi. B) Weeks 25-28 (IHC 100X): moderate stain in the vessel walls and strong stain in the stroma of stem villi; mild to moderate stain in the vessel walls and moderate to strong stain in the stroma of immature intermediate villi. Weak to mild stain in the vessel walls and a variable stain in the stroma of intermediate mature villi; weak to mild stain in the vessel walls and a variable (from weak to moderate) stain in the stroma of terminal villi. C) Weeks 29-32 (IHC 100X): variable (from mild to strong) stain in the vessel walls and in the stroma of stem villi and intermediate immature villi. Weak to moderate stain in the vessel walls and in the stroma of intermediate mature villi; weak to mild stain in the vessel walls and a weak to mild stain in the stroma of terminal villi. D) Weeks 33-36 (IHC 100X): mild to moderate stain in the vessel walls and moderate to strong stain in the stroma of stem villi; mild to moderate stain in the vessel walls and moderate to strong stain in the stroma of immature intermediate villi. Weak to mild stain in the vessel walls and in the stroma of intermediate mature villi; weak stain in the vessel walls and a weak to mild stain in the stroma of terminal villi.

### 3.1 Premature placentas

Each case we analysed showed positivity for staining, but with different intensity; no negative staining was observed. Figures 1 and 2 show an evident spatio-temporal variation in the staining intensity for both SMA and desmin in vascular wall and in stroma, with decreasing expressions not only during the pregnancy (temporal variation), but from proximal to distal part of the villous tree (spatial variation) being higher in the stem villi.

#### 3.1.1 Weeks 21-24 (n=3)

SMA positivity in stem villi was well defined and was evident both in vascular walls and stroma; vascular walls got strong expression of contractile elements, while stroma got moderate. Expression of intermediate immature villi was like stem villi. In the few intermediate mature villi, positivity was less marked than what was seen in stem villi, but was clearly expressed both in vascular walls and stroma. Vessel walls got moderate staining in all of cases, whereas stroma got mild expression. These results were found in all the analysed fields, it was not a focal finding (Table 2 and Figure 1a).

**Table 2:**
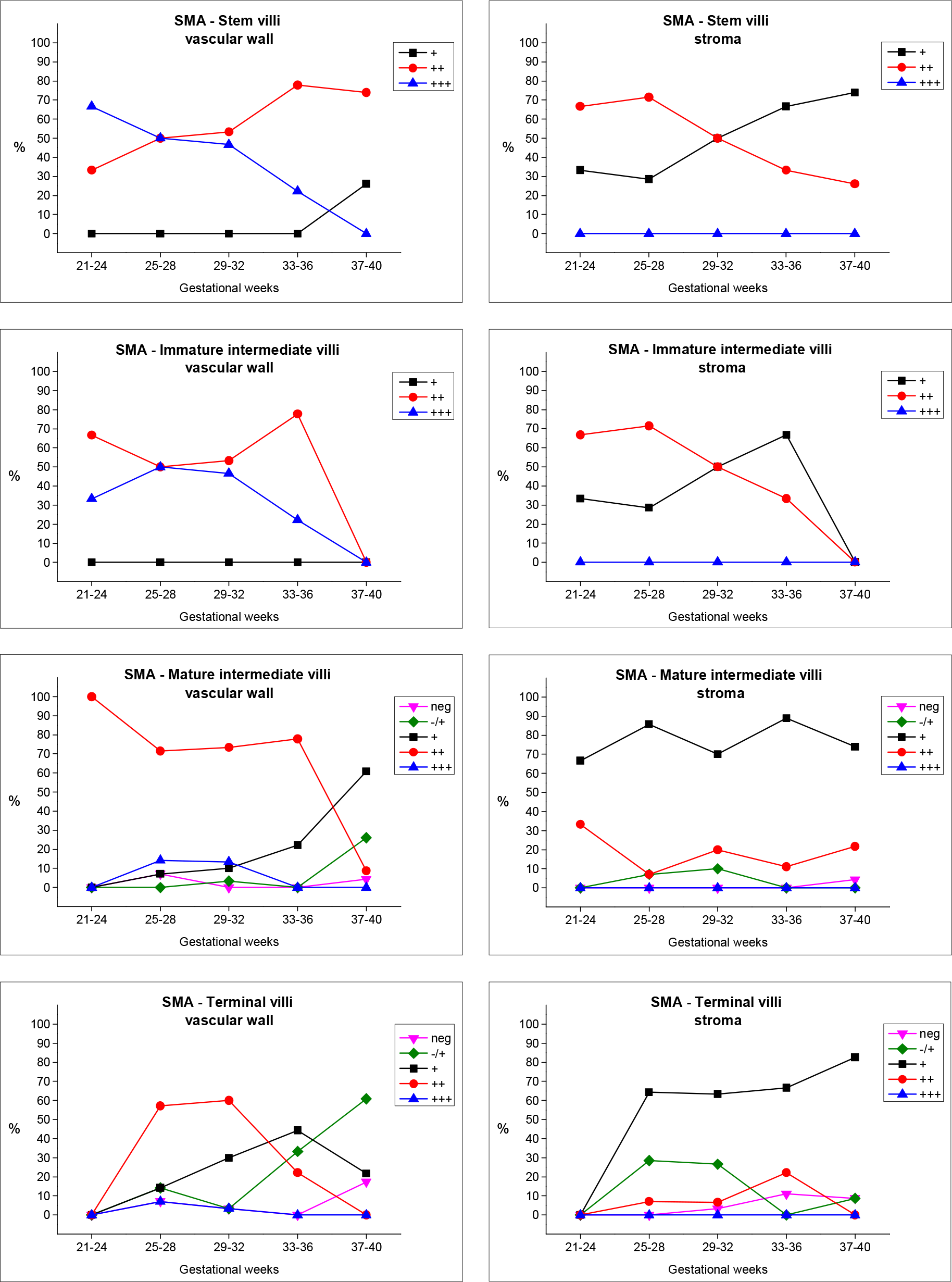
Temporal Variations of Smooth Muscle Actin (SMA) staining

Considering desmin IHC, vessel walls of stem villi and immature intermediate villi showed moderate staining, and their stroma stained strong in all of cases. 67% of mature intermediate villi cases presented mild expression in the vessel walls, whereas stroma stained moderate (Table 3 and Figure 2a).

**Table 3:**
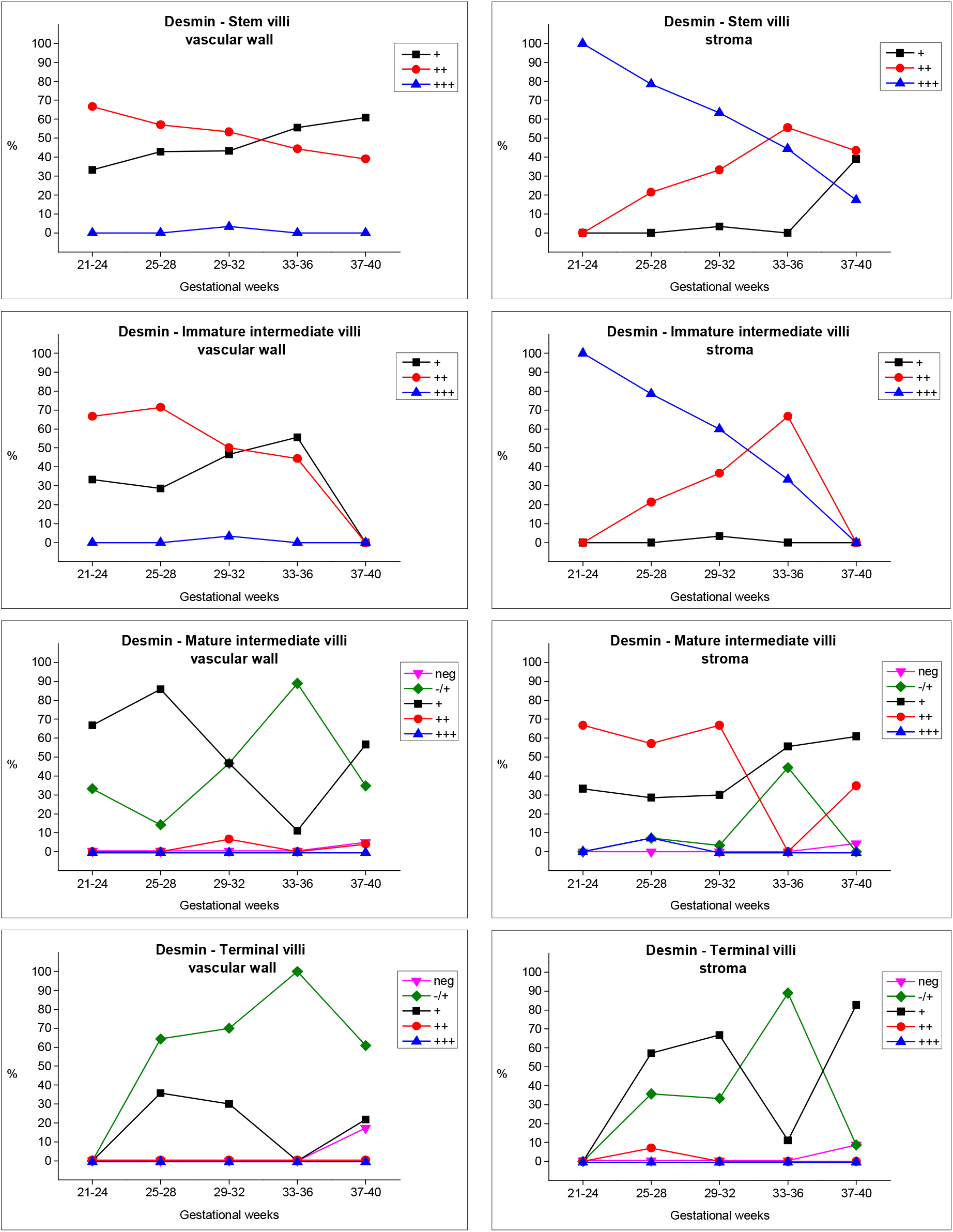
Temporal Variations of Desmin staining

#### 3.1.2. Weeks 25-28 (n=14; 3 twins)

SMA positivity in stem and intermediate immature villi was well defined and evident both in blood vessel walls and stroma. Blood vessel walls presented a moderate staining in half of cases and the other half had mild expression, whereas stroma presented strong and moderate expressions. While the presence of contractile elements in blood vessel walls of mature intermediate villi was mainly moderate, the stroma presented a mild staining. Positivity was also present in terminal villi, although to a far lesser extent than in other subtypes. Villous vessel walls showed moderate expression, but stroma showed mostly mild (Table 2 and Figure 1b).

The presence of contractile elements was also confirmed by positive staining with the anti-desmin antibody. In vascular walls of stem villi, the expression was mild and moderate, but stroma stained predominantly strong. Blood vessel walls of intermediate immature villi had moderate expression but stroma got strong. Blood vessel walls of mature intermediate villi had mild expression and stroma had mainly moderate. Vessel walls in terminal villi showed weak expression, whereas stroma stained mild (Table 3 and Figure 2b).

#### 3.1.3. Weeks 29-32 (n=30; 6 twins)

Vessel walls and stroma got SMA positivity, although slightly less evident than the placentas of earlier gestational weeks (21-28 gestational weeks). Blood vessel walls of stem and immature intermediate villi presented both moderate and strong expression in half of cases for each one, whereas stroma got mild and moderate staining with similar values. Blood vessel walls of mature intermediate villi had moderate staining, whereas stroma got mild. Positivity was present and evident in terminal villi; blood vessel walls had moderate expression and stroma had mild (Table 2 and Figure 1c).

Desmin IHC staining resulted like weeks 25-28 (Table 3 and Figure 2c). Blood vascular walls of stem villi presented mainly moderate staining, while stroma stained strong. Blood vessel walls of immature intermediate villi stained mild, but strong for stroma. Blood vessel walls of mature intermediate villi got both weak and mild desmin expression with 46.67%, while stroma stained moderate. Blood vessel walls of terminal villi stained weak (70%) and stroma stained mild (67%).

#### 3.1.4. Weeks 33-36 (n=9; 2 twins)

SMA positivity showed contractile elements both in vascular walls and stroma like observed in weeks 29-32, but slightly less evident than weeks 21-28. Blood vessel walls of stem villi stained moderate (77.78%) and strong (22.22%), whereas stroma stained mostly mild (67%). The extremely rare immature intermediate villi showed contractile elements in both stroma (mild expression) and vascular walls (moderate staining). Positivity in mature intermediate villi was well represented in the vascular walls like observed for stem villi, and stroma stained mild in 88.89% of cases. Positivity in terminal villi was evident but less marked than what was observed in the other villous subtypes; blood vessel and stroma presented mild expression (Table 2 and Figure 1d).

The expression of desmin was weaker than in earlier gestational weeks. Blood vascular walls of stem villi stained mostly mild and stroma moderate in 55.56% of cases for each one. Blood vessel walls of immature intermediate villi stained mild (55.56%) and stroma moderate (67%). Blood vessel walls of mature intermediate villi stained weak (88.89%) whereas stroma stained mild (55.56%). Blood vessel walls of terminal villi presented a weak expression in all the cases, but stroma stained weak only in 88.89% (Table 3 and Figure 2d).

### 3.2 Term placentas (n=23)

37th week: contractile elements were highlighted by staining for SMA in the vascular walls and in the stroma of stem and mature intermediate villi; in terminal villi the presence of SMA was especially noticeable in the stroma, but also in the vascular walls, albeit less clearly than other villous subtypes. Positivity for staining with anti-desmin antibody was like for SMA in stem and mature intermediate villi; it was expressed more mildly in terminal villi where it was especially positive in the stroma.

38th week: positive staining for SMA was highlighted in vascular walls and stroma of stem villi; the contractile component of mature intermediate villi was more evident, especially in the vascular walls but also in the stroma, as can was observed in terminal villi where positivity was present mainly in villous stroma. Positivity for anti-desmin antibody was also found, especially in vascular wall and stroma of stem villi; mature intermediate villi expressed staining predominantly in the stroma, whereas positivity was weaker and focal in the vascular walls of both these villi and terminal villi.

39th week: contractile elements were clearly present in the stroma and vascular walls of stem villi; SMA staining was also positive in the vascular walls and in the stroma of mature intermediate villi; terminal villi showed stronger positivity in the stroma and weaker positivity in the vascular walls. The anti-desmin antibody revealed the presence of contractile elements in the stroma and in the vascular walls of stem and mature intermediate villi; terminal villi showed positivity for this antibody especially in the stroma, although weaker than in other villous subtypes.

40th week: the expression of contractile elements, highlighted by SMA, was evident in the vessel walls and in the stroma of stem and mature intermediate villi; in terminal villi clear expression was present in the stroma, and in a weaker and more focal way in the vessels. Anti-desmin staining was like the 39th week, with evident positivity in the stroma and the vascular walls of the stem and mature intermediate villi; terminal villi expressed positivity in a weaker and focal way, especially in the stroma. Evident positive staining for SMA was also found in stem villi and even in mature intermediate villi of at-term placentas; in terminal villi the positivity was mainly expressed in the stroma and focally in the vascular walls as well, with varying modulation depending on the pathology. The presence of contractile elements was confirmed by anti-desmin antibody positivity, which was mainly present in the stroma and in the vascular walls of stem and mature intermediate villi.

## 4. Discussion

The fundamental morphologic works of Demir, Kohnen, and Kaufmann [1-4,9] stimulated our curiosity to evaluate a possible modulation of expression of the contractile component in the stroma and in the vessel walls of mature intermediate and terminal villi.

We chose to use SMA antibody since this isoform of actin allows to evaluate the myogenous differentiation [28]; this protein is seen in tissue with “pure” myogenic differentiation, but is also demonstrable in cells with myofibroblastic or myoepithelial features (smooth muscle cells in the media of vessels) and in pericytes [26]. We chose also desmin as a control antibody: desmin is a cytoplasmic intermediate filament protein characteristically found in muscle cells; in smooth muscle cells it is seen with cytoplasmic dense bodies and subplasmalemmal dense plaques. It’s often expressed in myogenous cells but also in fibroblasts, implicating a “myofibroblastic” nature for these cells and, along with alpha actin smooth muscle, is found also in pericytes [27,28].

Our findings revealed a difference in the intensity of staining: SMA was overall highly expressed than desmin both in preterm and in term placentas, both in vascular walls and in the stroma. However, both staining, also if singularly evaluated, were found in the same elements, such as smooth muscle cells in vascular walls, pericytes and myofibroblasts in the stroma, even if with different intensity in the various groups. An interesting result was observed in all premature subgroups: positivity for SMA persists in the stroma and in the vascular walls of mature intermediate villi and terminal villi, thus proving the presence and persistence of contractile elements in the vascular walls of these two subtypes of villi. Our work revealed the presence of leiomuscular-type contractile elements both in blood vessel walls and in the stroma of each type of villi (stem, immature intermediate, mature intermediate and terminal) and in each of the four groups we examined for high prematurity.

An extensive review of English literature from 1979 [1-28] was performed, focusing on villous blood vessels and stromal contractile component, both in at-term and in preterm placentas, as well as articles on immunohistochemical staining with anti-actin smooth muscle and anti-desmin antibodies. Only few of these articles describe preterm placentas [4,7,8,9], but from a pathological point of view, as in preeclampsia, IUGR or HELLP syndrome [5,14,25]; in other articles attention is focused on stem villi [2,3,15] allowing a diagnostic approach to vascular diseases of the placenta. Benirschke et al [9] compiled a table based on the works of Demir [2] and Kohnen [3], which is still considered the reference point for this issue: the table describes how immunohistochemical staining for desmin in at-term placentas shows positivity in all villous subtypes except for the undifferentiated mesenchymal villi. On the other hand, positivity for staining with anti-smooth muscle actin antibody is present in the stem villi of first, second and third degree, in the fibrous stroma as well as in the media of arterial and venous vessels; it is also present in the fibrous part around the larger vessels of immature intermediate villi. In contrast, a lack of response to this antibody is observed in both mature intermediate and terminal villi.

We focused our attention on the terminal parts of the villous tree, where maternal-foetal exchange takes place, to evaluate the possible maturational alterations of the villous tree itself in high prematurity stages, based on the adaptation or the pathological conditions that cause high prematurity. Even more unexpectedly, in at-term placenta control cases the presence of smooth muscle cells was observed in all subtypes of villi, but mainly in the most peripheral ramifications. Most of the articles discussing at-term placentas analysed only stem and immature intermediate villi [7,8] or described them as having contractile elements [2,3]. Our results are similar both in preterm and at-term placentas, indicating that the contractile component should be evaluated in terms of modulation or typology of expression, but not as an absence/presence binomial, even in at-term placentas. In our opinion, the presence of contractile elements, not so much in the stroma and in the vascular walls of stem villi (which is well-established in the literature) as in mature intermediate and terminal villi, changes the diagnostic approach to these placentas. The modulating and dynamic concept of contractile activity of both the vessel walls and stroma of villi is introduced, and this is to be correlated with the pathological patterns of the placenta, or with clinical conditions of the mother or in any case with the dynamics of the maternal-foetal exchange.

## 5. Conclusions

Based on our analysis and review of the literature, which lack of descriptions of these villi, we may conclude that high premature placentas have a contractile component within each type of villi that is made up of leiomuscular cells or myofibroblasts which are highlighted by staining with anti-smooth muscle actin and desmin antibodies. These components may be present both in the vascular walls and in the stroma of villi, but with different expression of staining intensity, depending on the possible pathology affecting the placenta and/or the mother.

Mature intermediate and terminal villi have a muscular component even in term placentas, with a degree of staining that changes depending on the pathology. There may be an increase in staining in cases of preeclampsia (especially in the vascular walls), a condition in which the resistance of peripheral vessels of the placenta increases. Another common condition is diabetes, where there is a lower presence of staining for actin, especially in the stroma of villi. In highly premature placentas, four groups were identified based on the progressive complication of villous ramifications and on the appearance of exchanging terminal villi. This means that placental exchange changes progressively from a generic exchange in every vase of every order of villi to a specific exchange in mature intermediate and terminal villi, because the exchanging process needs to become maximum. To imagine that these terminal villi do not have in the stroma (as one can find in the literature) or also in the vascular walls (as one can comprehend from literature but without a clear description) a contractile component, means giving these vases a passive hemodynamic connotation, so an idea of a static nature. On the contrary, finding muscular components means that mature intermediate and terminal villi, which appear progressively in various groups and stages, are also active in the modulation of the blood flow. This imposes a revisitation not only of the physiopathology of the placental circulation but also of the possible implications for the target of pharmacological therapies.

## Author Contributions

VGV, MP, RB and EF conceived and designed the study, and wrote, edited and reviewed the manuscript. GS, CMB, FaB, DB and FrB researched and analysed data, and wrote, edited and reviewed the manuscript. All authors gave final approval for publication. VGV and EF take full responsibility for the work, including the study design, access to data and the decision to submit and publish the manuscript.

## Acknowledgments

Thanks to Leonardo Penuela MD PhD for his support in editing. Special Thanks to Mrs Giorgia Anselmi for her technical support.

